# Suppressing phagocyte activation by overexpressing the phosphatidylserine lipase ABHD12 preserves sarmopathic nerves

**DOI:** 10.1101/2024.06.20.599919

**Authors:** Caitlin B. Dingwall, Yo Sasaki, Amy Strickland, Daniel W. Summers, A. Joseph Bloom, Aaron DiAntonio, Jeffrey Milbrandt

## Abstract

Programmed axon degeneration (AxD) is a key feature of many neurodegenerative diseases. In healthy axons, the axon survival factor NMNAT2 inhibits SARM1, the central executioner of AxD, preventing it from initiating the rapid local NAD+ depletion and metabolic catastrophe that precipitates axon destruction. Because these components of the AxD pathway act within neurons, it was also assumed that the timetable of AxD was set strictly by a cell-intrinsic mechanism independent of neuron-extrinsic processes later activated by axon fragmentation. However, using a rare human disease model of neuropathy caused by hypomorphic NMNAT2 mutations and chronic SARM1 activation (sarmopathy), we demonstrated that neuronal SARM1 can initiate macrophage-mediated axon elimination long before stressed-but-viable axons would otherwise succumb to cell-intrinsic metabolic failure. Investigating potential SARM1-dependent signals that mediate macrophage recognition and/or engulfment of stressed-but-viable axons, we found that chronic SARM1 activation triggers axonal blebbing and dysregulation of phosphatidylserine (PS), a potent phagocyte immunomodulatory molecule. Neuronal expression of the phosphatidylserine lipase ABDH12 suppresses nerve macrophage activation, preserves motor axon integrity, and rescues motor function in this chronic sarmopathy model. We conclude that PS dysregulation is an early SARM1-dependent axonal stress signal, and that blockade of phagocytic recognition and engulfment of stressed-but-viable axons could be an attractive therapeutic target for management of neurological disorders involving SARM1 activation.

## Introduction

Neuroinflammation is a double-edged sword^1^. In the CNS, microglia are the primary resident phagocytes comprising a highly dynamic pool of cells involved in both neuroprotective and neurotoxic functions^2^. Phagocytes mediate debris clearance and tissue repair^3^, but sustained inflammation can also drive the progression of neurodegeneration and is a frequent feature of disorders including AD, PD, FTD, ALS, and peripheral neuropathies^4–6^. Targeting neuroinflammation in preclinical models of these diseases has shown symptomatic reversal^7^, suggesting that neuroinflammation may worsen neuropathology. Moreover, functional recovery in these models suggests that immunocytes target viable neurons, not only those that are already irreversibly fated for cell death. Understanding the signals that govern phagocytic recognition and engulfment of compromised neurons is essential to designing precision immunomodulatory therapeutics for treating neurodegenerative disease. Our recent work has shown that SARM1, the central executioner of axon-intrinsic degeneration, participates in coordinating phagocytic axon loss in the peripheral nervous system^8^.

Programmed axon degeneration (AxD) is an active, genetically-encoded subcellular destruction pathway^9^, conceptually akin to apoptosis. In a healthy neuron, axon integrity is maintained by opposing actions of two enzymes: an NAD+ synthetase, NMNAT2^10,11^, and an NAD+ hydrolase, SARM1^12^. SARM1 is normally kept inactive by the presence of NMNAT2 in the axon^13–16^. In a prior study, we created the first mouse model of chronic SARM1 activation (termed “sarmopathy”) based upon rare *Nmnat2* mutations identified in two patients with a devastating progressive neuropathy syndrome^8^. Macrophage depletion using CSF1 receptor antibody blocks axon loss in this model, suggesting that neuronal SARM1 also promotes non-cell autonomous axon degeneration via a mechanism requiring macrophages. Moreover, the persistence of sarmopathic motor axons and functional recovery after macrophage depletion supports the hypothesis that macrophages target stressed-but-viable axons. These results raised a fundamental question: how does chronic SARM1 activity within axons mediate the changes in macrophage behavior that drive degeneration in sarmopathic peripheral nerves?

One candidate signal classically involved in macrophage recognition and engulfment is phosphatidylserine (PS), the quintessential ‘‘eat me’’ molecule exposed on the surface of apoptotic and metabolically compromised cells^17–19^, including during axon degeneration^20^. SARM1 activation drives calcium influx in damaged axons, an event that precedes the axonal membrane externalization of PS^21^. SARM1 is required for PS exposure in models of axotomy and vincristine-induced axon degeneration^22^. *Drosophila* studies demonstrated that reducing PS exposure on degenerating neurites can abrogate the phagocyte response and slow neurite loss^23^. Conversely, ectopic PS exposure on otherwise healthy neurites triggers phagocyte-mediated neurite breakdown^23^. Moreover, under pathologic conditions, such as during cellular stress, levels of the bioactive PS derivatives oxidized PS (oxPS) and lysoPS rise and subsequently trigger robust immune activation^24–30^. Both oxPS and lysoPS are degraded by the PS lipase ABHD12^31^ and patients with *ABHD12* loss-of-function mutations develop PHARC syndrome, a rare congenital neurodegenerative disease involving pro-inflammatory lysoPS and oxPS accumulation in the brain, enhanced microglial phagocytosis, and pathological neuroinflammation^32^. These data suggest that glial or immunocyte recognition of PS could drive phagocytic clearance of stressed-but-viable axons that might otherwise persist and potentially recover. By contrast, strictly cell-autonomous axon degeneration may be mainly a phenomenon of severe traumatic injury to clear irreversibly compromised axons e.g. classical Wallerian degeneration following axon transection.

To test fundamental mechanisms governing sarmopathic axon loss, we generated an in vitro system to model chronic SARM1 activation in cultured sensory neurons. Here, we show that in vitro chronic SARM1 activation absent acute axon injury triggers slow axon degeneration rather than spontaneous axon fragmentation, supporting our observation from the sarmopathy mouse model that axon loss is not an all-or-nothing event in chronic SARM1-driven disease. We found that sarmopathic axons develop axonal blebs and expose phosphatidylserine well before frank axon degeneration occurs. In addition, acute activation of SARM1 in cultured neurons provokes a robust rise in the pro-inflammatory PS species lyso-phosphatidylserine (lysoPS) which can be blocked by SARM1 deletion. ABHD12 function has been primarily studied in microglia and macrophages, however here we demonstrate that inhibition of ABHD12 in neurons is sufficient to trigger a robust increase in lysoPS, demonstrating a heretofore unappreciated function in neurons. Since ABHD12 functions in neurons, we engineered an AAV-mediated in vivo gene therapy strategy to overexpress ABHD12 in neurons and thus lower levels of pathologic PS. Excitingly, ABHD12 overexpression abrogated nerve macrophage activation in our sarmopathy model without reducing total macrophage numbers, suggesting that pathologic PS regulates macrophage function rather than recruitment/proliferation. In addition, reduced macrophage activation correlated with preservation of femoral nerve axons and motor function, suggesting that sarmopathic axons targeted for removal are indeed still viable. Our study demonstrates that PS exposure is a key step in the *in vivo* axon degeneration pathway and that pathologic PS is a critical early axonal targeting signal involved in sarmopathic disease pathogenesis. Collectively, our data suggest that suppressing this early distress signal may allow persistence or recovery of stressed-but-viable axons and have broad therapeutic value to treat neurodegenerative diseases involving chronic SARM1 activation, including ALS^33,34^ and peripheral neuropathies^8,35–39^.

## Results

### Absence of NMNAT2 is sufficient to evoke chronic SARM1 NADase activity in uninjured cultured sensory neurons

The prevailing evidence demonstrates that acute NMNAT2 loss triggers a rise in the NMN/NAD+ ratio leading to SARM1 activation^40–43^. This observation would predict that NMNAT2 loss or deficiency would trigger acute SARM1-mediated axon degeneration. However, in our sarmopathy model created using two hypomorphic *Nmnat2* variants, we find that SARM1 is chronically activated leading to a slowly progressive neuropathy; thus, axon loss does not occur as an all-or-nothing event^8^.

Genetic deletion of *Nmnat2* alone is lethal due to failed axon elongation during development^44,45^. However, mice lacking both NMNAT2 and SARM1 (NMNAT2/SARM1 dKO mice) are viable^16^, providing a powerful background to interrogate the destructive pathway, including differences between injury-evoked axon degeneration versus genetic NMNAT2 loss. We cultured dKO dorsal root ganglion (DRG) neurons, allowed their axons to grow and mature, then reintroduced a *SARM1* transgene to test the effects of SARM1 in the absence of NMNAT2 in an otherwise healthy neuron. At baseline, intact dKO axons exhibit decreased NAD+ and ATP due to a lack of NMNAT2 but remain otherwise morphologically normal (**Figure 1A**). Prior studies show that when SARM1 is re-expressed in SARM1 KO neurons, they display wild-type levels of NAD^+^, ATP, and NAD^+^ consumption, low levels of cADPR (a SARM1-specific NAD metabolite^46^), and normal axon degeneration in response to injury^15,47^. We examined axonal metabolites three days after SARM1 lentiviral transduction and found that SARM1 addback to dKO neurons (dKO + SARM1) resulted in decreased NAD^+^ and ATP, elevated cADPR, and an increased rate of NAD^+^ consumption, demonstrating that SARM1 is activated (**Figure 1B-C**). The increase in NAD+ consumption observed in dKO + SARM1 axons was blocked by co-expression of NMNAT2, confirming the current paradigm that NMNAT2 suppresses SARM1 activation (**Figure 1C**).

**Figure 1.**
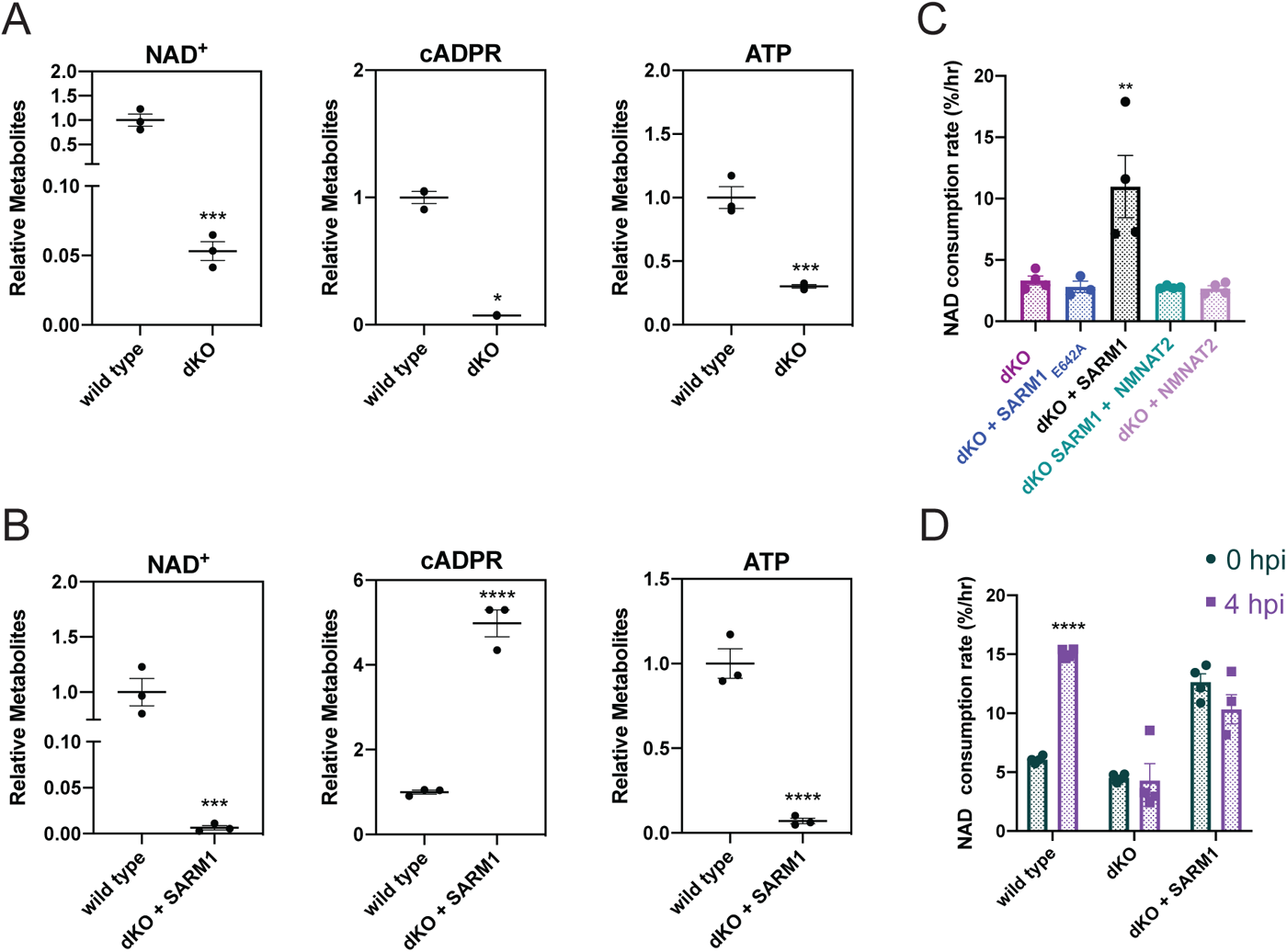
Genetic loss of NMNAT2 is sufficient to drive SARM1 activation in the absence of injury. **(A-B)** Axonal metabolites from DIV8 WT, dKO (control), and dKO + SARM1 neurons (n=3). **(C)** NAD+ flux assay on DIV8 WT or dKO axons (control, wild-type SARM1, SARM1_E642A_, or wild-type SARM1 + NMNAT2) (n=4). **(D)** NAD+ flux assay on DIV8 WT, dKO (control), and dKO + SARM1 axons 4 hours after injury (n=4). All data are presented as mean ± SEM. Statistical significance determined by Student’s unpaired t-test. ns: not significant, *p<0.05, **p<0.01, ***p<0.001, ****p<0.0001.

To date, most in vitro axon degeneration studies have employed acute axon injury paradigms (i.e. axotomy) which induces rapid NMNAT2 loss in addition to the other effects of acute injury. To establish whether the level of SARM1 activation is equivalent between sarmopathy caused by genetic NMNAT2 loss and axotomy-induced sarmopathy, we investigated whether injury enhances SARM1 NADase activity independent of NMNAT2 loss. In injured wild-type sensory neurons, SARM1 activation leads to increased NAD+ consumption most evident 4 hours after insult^15^. Since re-expression of SARM1 in dKO neurons and injuring wild-type neurons yield similar rates of NAD+ consumption, we hypothesized that both conditions already induce maximum SARM1 activation and therefore injury of dKO + SARM1 axons would not further enhance SARM1 NADase activity. To test this, we axotomized both wild-type and dKO + SARM1 sensory neurons and performed an axonal NAD+ flux assay to measure relative NAD+ consumption. This confirmed that NAD+ consumption in uninjured dKO + SARM1 axons was not significantly different than injured wild-type axons (**Figure 1D**). Furthermore, injured dKO + SARM1 axons did not exhibit increased NAD+ consumption compared to intact dKO + SARM1 axons (**Figure 1D**). These findings demonstrate that absence of NMNAT2 is sufficient to evoke robust SARM1 NADase activity that is not further enhanced by injury. Given that SARM1 is chronically activated in the absence of NMNAT2, we henceforth term axons with deficient NMNAT2 activity “sarmopathic”.

### SARM1 activation induces axonal blebbing in uninjured neurons

We next tested whether our in vitro dKO sarmopathic axons would degenerate akin to what has been observed in axotomy-induced axon degeneration. We cultured dKO DRG sensory neurons for five days in vitro (DIV) before reintroducing SARM1 using lentiviral transduction (**Figure 2A**). As a control, we also used the enzymatically-dead SARM1^E642A^ mutant, which is unable to trigger axon degeneration^12^. Axonal health was monitored over the next four days using our automated image analysis axon degeneration assay which scores degeneration based on axon fragmentation. Strikingly, despite being fully active, reintroduction of wild-type SARM1 was not sufficient to induce spontaneous axon fragmentation; instead, axons developed prominent membrane blebs (**Figure 2B**). In contrast, expression of SARM1_E642A_ had no effect on axon morphology (**Figure 2C**) demonstrating that SARM1’s NADase activity is required for the formation of axonal blebs. The blebs developed between the second and third day of wild-type SARM1 re-expression and persisted for the duration of the experiment without progressing to axon degeneration (**Figure 2C**). Surprisingly, transection of these “blebbing” dKO + SARM1 axons did induce complete axon fragmentation, suggesting that injury stimulates a second, pro-degenerative pathway or triggers loss of a second axon maintenance pathway that is required to progress from axon blebbing to fragmentation. (**Figure 2B, D**). These findings reveal a previously unappreciated NMNAT2-independent step in SARM1-evoked axon degeneration.

**Figure 2.**
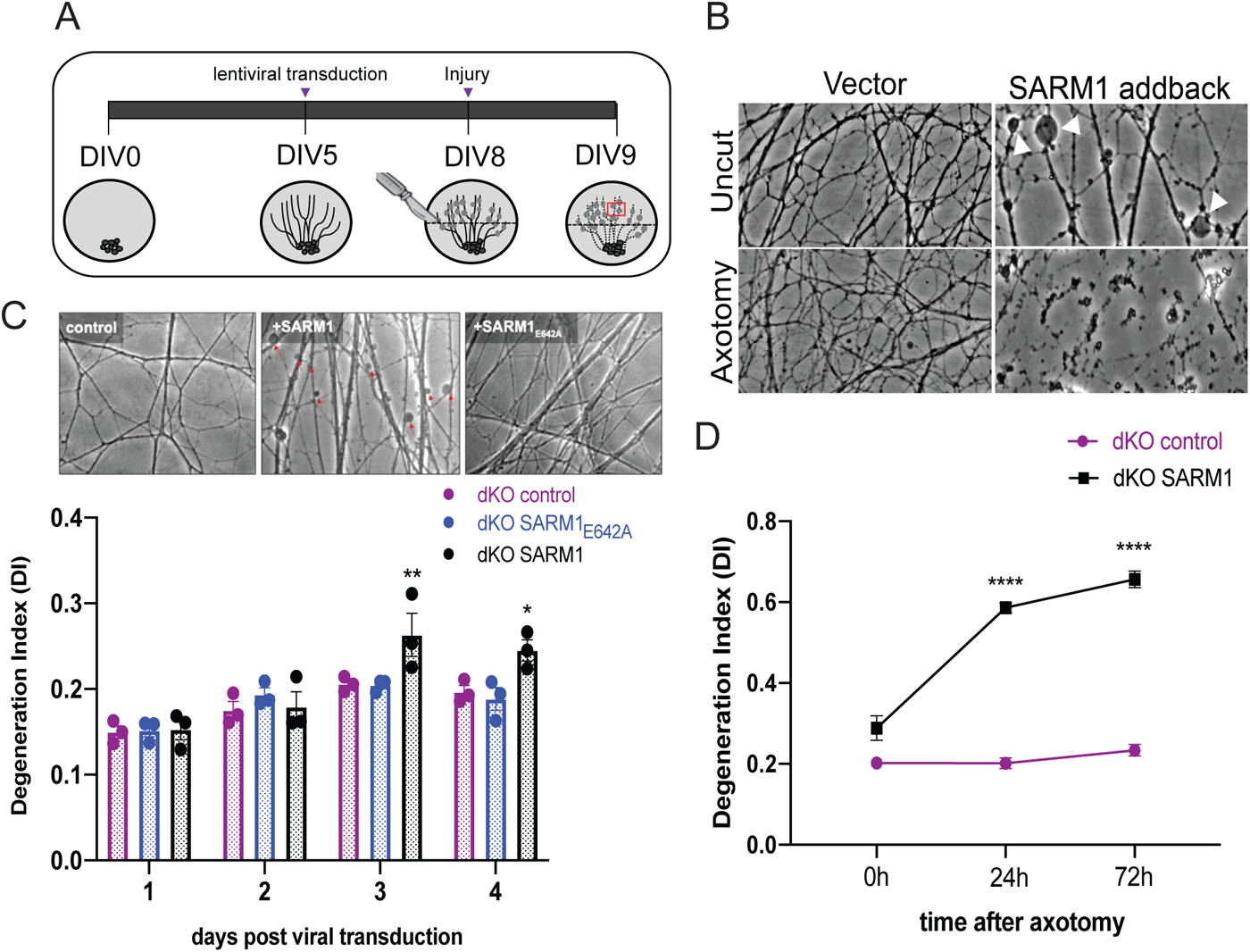
SARM1 activation induces uninjured axons to bleb but not fragment. **(A)** Schematic of lentiviral transduction paradigm. **(B)** Representative images of uncut and axotomized axons of dKO (control) and dKO + wild-type SARM1 neurons. (**C)** Representative images of uninjured dKO (control), dKO + wild-type SARM1, and dKO + SARM1_E642A_ axons on DIV8. Quantification of axon degeneration post-lentiviral transduction in dKO neurons using the degeneration index (DI). Y-axis: 0.0 indicates morphologically intact axons and 1.0 indicates complete axon fragmentation, max Y-axis value displayed is 0.4 (n=3). (**D)** Quantification of axon degeneration after axotomy of dKO (control) and dKO+SARM1 axons using the degeneration index (DI). (n=3). All data are presented as mean ± SEM. Statistical significance determined by Student’s unpaired t-test. ns: not significant, *p<0.05, **p<0.01, ***p<0.001, ****p<0.0001.

### SARM1 activation triggers membrane PS exposure on stressed-but-viable axons

Following axonal injury in wild-type neurons, there is an early “latent” period of programmed degeneration that lasts 4-6 hours characterized morphologically by the emergence of axonal blebbing culminating in axon fragmentation^21,48^. Similarly, treating wild-type axons with the mitochondrial poison rotenone triggers SARM1 activation and precipitates axonal blebbing akin to that seen in traumatic axonal injury^49^. Rotenone treated axons also recover both metabolically and morphologically— i.e. axon blebbing is reversed—after acute pharmacologic SARM1 inhibition^49^. Such a reversal suggests that blebbing axons are metabolically stressed but still viable. During this window of latent axon degeneration, several subcellular events occur, including phosphatidylserine exposure on the axonal membrane^21^.

The phospholipid phosphatidylserine (PS), which is normally present primarily in the inner leaflet of the plasma membrane and excluded from the outer leaflet of healthy neurons^50^, is exposed on the external leaflet of the axon membrane during bioenergetic stress^21^. Moreover, exposed PS is concentrated on axonal swellings^22,51^. Up until this point, nearly all studies on PS exposure on compromised axons have been done in acute injury conditions^21–23,51^. Here we leveraged our dKO model to examine whether PS exposure also occurs with chronic SARM1 activation. PS exposure is detected using the cell-impermeable PS ligand Annexin V^28^. We find that sarmopathic dKO + SARM1 axons display significant Annexin V staining, particularly on axonal blebs, indicating that PS is externalized to their extracellular leaflet. By contrast, relatively little Annexin V binds to healthy control axons (**Figure 3A-C**). Altogether, these findings suggest that chronically sarmopathic axons persist in a state that parallels the early phase of degeneration in injured axons.

**Figure 3.**
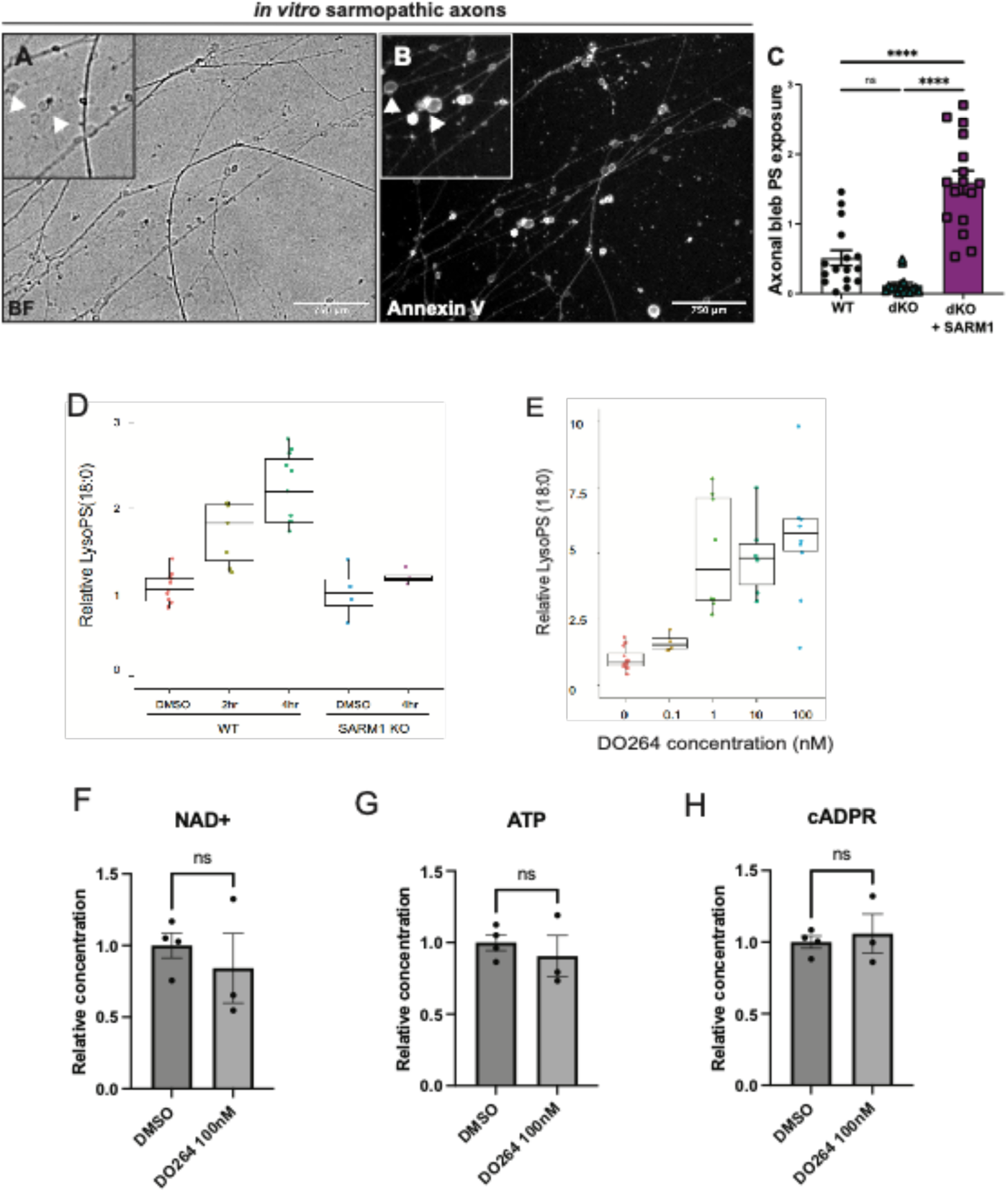
SARM1 activation triggers PS dysregulation in stressed-but-viable axons. **(A)** Representative (a) brightfield and (b) Annexin V fluorescent images of blebs on dKO axons transduced with SARM1. Insets depict magnified representative images of axonal blebs (arrows) and Annexin V signal on sarmopathic axons (dKO + SARM1). **(C)** Quantification of Annexin V intensity on axonal blebs from wild-type, dKO, and dKO + SARM1 neurons (n=16). **(D)** Quantification of relative lyso-PS levels as measured by LC-MS/MS after treatment of wild-type or SARM1 KO neurons with DMSO (control) or 50mM Vacor at 0, 2, and 4hrs. **(E)** Quantification of relative lyso-PS levels as measured by LC-MS/MS after 24hr treatment of wild-type neurons with increasing doses of the ABHD12 inhibitor DO264 (n=3). **(F)** Quantification of NAD+ levels after 48hr treatment of wild-type neurons with 100nM DO264 (n=3). **(G)** Quantification of ATP levels after 48hr treatment of wild-type neurons with 100nM DO264 (n=3). **(H)** Quantification of cADPR levels after 48hr treatment of wild-type neurons with 100nM DO264 (n=3). All data are presented as mean ± SEM. Statistical significance determined by Student’s unpaired t-test or one-way ANOVA with multiple comparisons. ns: not significant, *p<0.05, **p<0.01, ***p<0.001, ****p<0.0001.

Dysregulation of PS dynamics results in both the exposure of PS on the extracellular leaflet of the axonal membrane and the buildup of potent bioactive PS derivatives. Altered cellular bioenergetics can lead to excessive production of reactive oxygen species (ROS). Surplus ROS can react with lipid membranes^52^ such as PS, generating oxidized PS (oxPS) that is readily externalized to the outer leaflet of the PM where it acts as an immunomodulatory signal^29,30^. In addition to the production of oxPS, another potent bioactive PS derivative, lysophosphatidylserine (lyso-PS), regulates several immunologic processes including macrophage activation^24–26,53^. To examine whether oxPS and lysoPS levels are altered by SARM1 activation, we treated cultured wild-type and *Sarm1* KO DRG sensory neurons with the potent SARM1 activator Vacor^54^. Due to technical limitations, we were unable to measure oxPS; however, within four hours of treatment, we observed an over two-fold increase in lysoPS levels in wild-type neurons but not in *Sarm1* KO, demonstrating that SARM1 activation induces lysoPS production (**Figure 3D**).

ABHD12, a PS phospholipase, regulates levels of several potent pro-phagocytic PS derivatives including both oxPS and lysoPS. A murine *Abhd12* KO model develops massive CNS neuroinflammation^55^, oxPS and lysoPS accumulation, and increased phagocytic activity in microglia^56^. Human *ABHD12* mutations cause the early onset neurodegenerative disease PHARC (polyneuropathy, hearing loss, ataxia, retinitis pigmentosa, and cataracts)^57–61^. ABHD12 is thought to function primarily in microglia and macrophages to regulate PS metabolism^57^ and mediate inflammation and phagocytosis. To test whether ABHD12 similarly functions to regulate lysoPS metabolism in neurons, we treated cultured DRG sensory neurons with the specific ABHD12 inhibitor DO264^62^. Remarkably, inhibition of ABHD12 robustly increases cellular lysoPS in otherwise healthy neurons (**Figure 3E**), strongly suggesting that ABHD12 is an endogenous lysoPS regulator in neurons, contrary to the long-held belief that it acts principally in phagocytes. Critically, this rise in lysoPS did not induce changes in NAD+, ATP, or the SARM1 biomarker cADPR^46^ (**Figure 3F-H**), confirming that lysoPS generation in this paradigm occurred due to pharmacologic inhibition of endogenous ABHD12 rather than off-target SARM1 activation.

### Axon blebbing occurs in a sarmopathy mouse model

Sarmopathy mice develop robust SARM1-dependent peripheral neuroinflammation early in disease, evidenced by abundant activated (CD68+) nerve macrophages and elevated total nerve macrophages as early as 2 months of age^8^. Depletion of tissue macrophages during the initial phase of disease prevented the neuropathy syndrome and preserved axons, suggesting that macrophages promote the neuropathology^8^.

We next examined whether this model exhibits morphologic and subcellular changes like those observed in cultured sarmopathic neurons. We intrathecally injected an EGFP construct (1.6X10^13^ virions) under the control of the synapsin promotor to fluorescently label peripheral nerve axons of young (3-month-old) *Nmnat2^V98M/R232Q^* sarmopathic mice. Sparse fluorescent labeling of axons in the femoral nerve revealed two morphologically distinct populations: axons with a consistent diameter and axons with prominent blebs akin to the sarmopathic dKO axons observed in vitro (**Figure 4A-A ’**). To examine macrophage-axon interactions, we immunostained the fluorescently labeled femoral axons with the activated macrophage marker CD68 and found that CD68+ macrophages reside closer to blebbing axons on average than to intact axons (**Figure 4B-D**) suggesting that macrophages are selectively attracted to blebbing axons prior to their frank degeneration.

**Figure 4.**
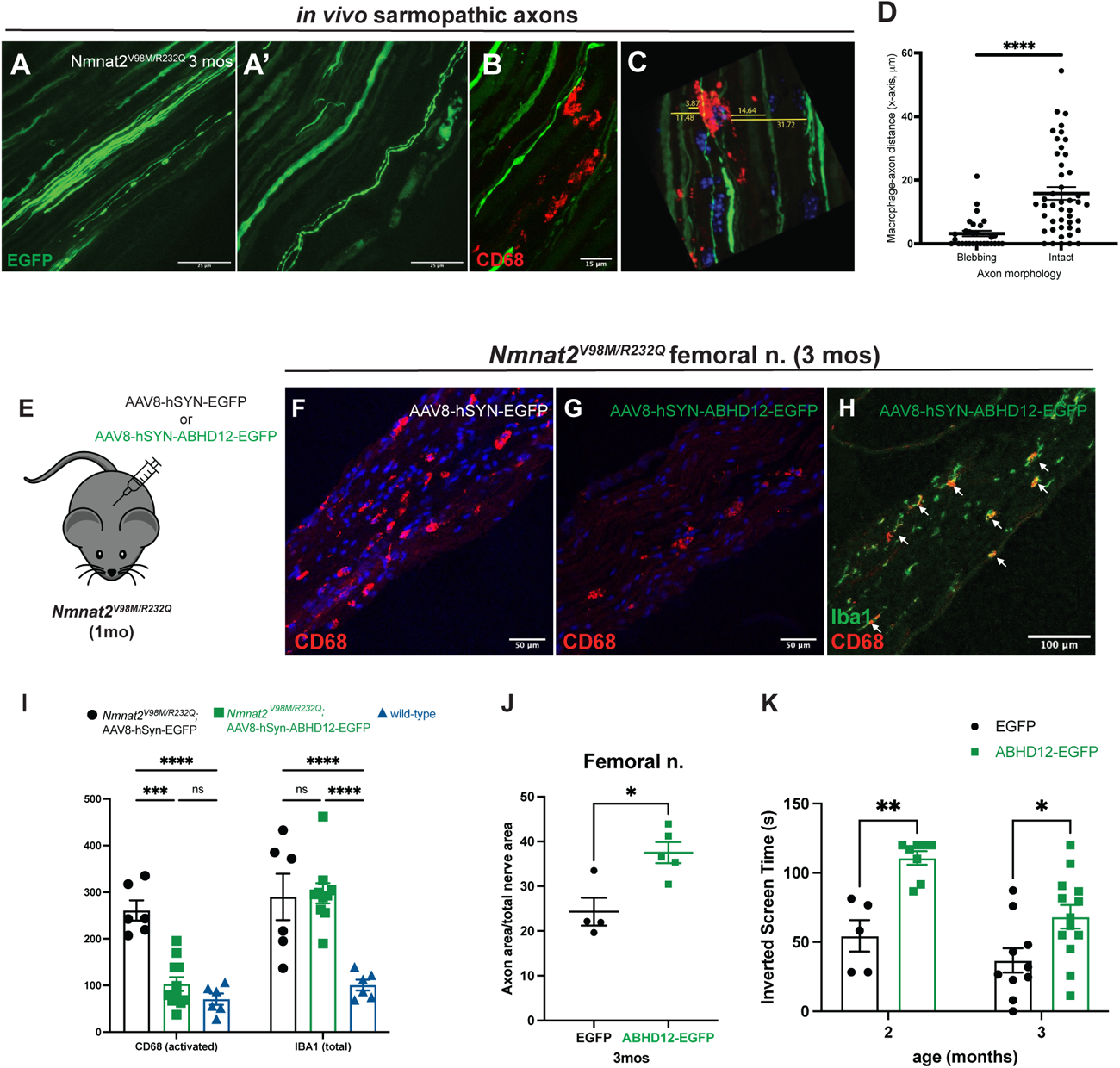
In vivo neuronal ABHD12 overexpression in sarmopathy mouse model reduces macrophage activation and rescues motor axon loss and neuropathy. **(A-A’)** Representative images of AAV-mediated, neuron-specific expression of EGFP in the distal femoral nerve of 3-month-old *Nmnat2^V98M/R232Q^* sarmopathy mice. EGFP expression reveals a population of blebbing axons. **(B)** Representative images of CD68 immunostaining and EGFP-labeled axons in femoral nerve of 3-month-old *Nmnat2^V98M/R232Q^*sarmopathy mice. **(C)** Schematic for macrophage-axon association quantification. **(D)** Macrophage-axon distance quantification in femoral nerve of 3-month-old *Nmnat2^V98M/R232Q^* sarmopathy mice (n=3 animals, each point replicates a counted axon). **(E)** Schematic of AAV-mediated, neuron-specific expression of EGFP or ABHD12-EGFP in 1-month-old *Nmnat2^V98M/R232Q^*sarmopathy mice. **(F-G)** Representative images of CD68 immunostaining in femoral nerve of 3-month-old *Nmnat2^V98M/R232Q^* sarmopathy mice after two months of control (EGFP) or *Abhd12* gene therapy. **(H)** Representative images of Iba1 and CD68 immunostaining in femoral nerve of 3-month-old *Nmnat2^V98M/R232Q^* sarmopathy mice after two months of *Abhd12* gene therapy. **(I)** Quantification of activated CD68+ and total Iba1+ macrophages in femoral nerve of 3-month-old *Nmnat2^V98M/R232Q^*sarmopathy mice after two months of control (EGFP) or *Abhd12* gene therapy. **(J)** Percent axonal area/total nerve area for femoral nerve calculated at 3mos (2mos of AAV-mediated gene therapy with control (n=4) or *Abhd12* IgG (n=5). **(K)** Average time suspended from an inverted screen (max. 120 seconds) for EGFP control (*n* = 10) and ABHD12 gene therapy *Nmnat2^V98M/R232Q^* (*n* = 14) mice at 2 and 3 months of age. All data are presented as mean ± SEM. Statistical significance determined by Student’s unpaired t-test or one-way ANOVA with multiple comparisons. ns: not significant, *p<0.05, **p<0.01, ***p<0.001, ****p<0.0001.

### In vivo neuronal ABHD12 overexpression in sarmopathy mouse model reduces macrophage activation and rescues motor axon loss and neuropathy

Given our finding that SARM1 activation triggers a rise in lyso-PS and membrane-exposed PS, we sought to investigate the role of PS in mediating sarmopathy. Our in vitro studies demonstrated that ABHD12 is an endogenous regulator of lyso-PS metabolism in neurons. Thus, to lower cellular levels of these pathologic PS species, we intrathecally injected one-month-old sarmopathy mice with AAV virions (1.3 x 10^13^) expressing ABHD12-EGFP or EGFP alone (control) driven by the neuron-specific synapsin promoter (**Figure 4E**).

Our prior work showed that sarmopathy mice develop a profound motor neuropathy and massive nerve macrophage infiltration and activation, both which are evident by two months of age. We first tested whether ABHD12 gene therapy reduces macrophage recruitment and found that the total number of nerve macrophages (identified by Iba1) did not significantly differ between control and ABHD12 gene therapy-treated nerves, remaining nearly three-fold higher than wild-type animals (**Figure 4H-I**), Nearly all nerve macrophages express the marker Iba1^63^, whereas only activated macrophages express CD68^64^. Strikingly, we found that the number of activated (CD68+) nerve macrophages in sarmopathy mouse femoral nerves was markedly decreased by the ABHD12 gene therapy, to levels comparable to that of wild-type animals (**Figure 4F-I, K**). Altogether, these data demonstrate that while ABHD12 gene therapy does not have a demonstrable effect on total macrophage recruitment and/or proliferation, it does lead to a profound inhibition of macrophage activation.

Prolonged macrophage depletion in sarmopathy mice rescues motor axon loss and motor dysfunction^8^. Thus, we next asked whether the reduced nerve macrophage activation after ABHD12 gene therapy would similarly lead to improvement in nerve pathology and motor function. By around 3 months of age, sarmopathy mice develop appreciable axon loss in the femoral nerve^8^, demonstrated as a reduction in total axon area in the nerve. To examine this, we performed microscopic analysis on toluidine blue-stained femoral nerves from 3-month-old sarmopathy mice and found motor axon loss was reduced in mice that received ABHD12 gene therapy compared to (EGFP) controls (**Figure 4J**). However, this analysis could not distinguish whether the additional axons were truly viable or merely remnants of dysfunctional axons prior to phagocytosis. Therefore, we assayed the motor function of ABHD12 gene therapy-treated sarmopathy mice using the inverted screen assay and found that ABHD12 gene-therapy significantly rescued their motor deficits (**Figure 4K**), akin to the therapeutic effect we observed after macrophage depletion^8^. Together, these data suggest that buildup of inflammatory PS derivatives contributes to nerve macrophage activation and loss of stressed-but-viable motor axons, and that gene therapies targeting PS dynamics after nerve perturbation are feasible and could ameliorate sarmopathy-associated motor decline.

## Discussion

Our recent findings demonstrated that stressed-but-viable axons are targeted for clearance by inflammatory cells long before they degenerate via strictly neuron-autonomous mechanisms^8,65^. Thus, contrary to the classic neuron-exclusive view, chronic SARM1 activation also precipitates a coordinated *transcellular* injury response to facilitate axon clearance. Here, we examined PS dysregulation as a SARM1-dependent early signal of compromised axons. Our work identified ABHD12 as an endogenous PS regulator in neurons and revealed that in vitro SARM1 activation triggers buildup of the pro-inflammatory PS derivative lyso-PS, PS externalization, and axonal blebbing. In our in vivo sarmopathy model we observed axonal blebbing and activated macrophages associated with pathogenic axons, highlighting a conserved program of SARM1-dependent axonal changes and neuroimmune signaling activation early in development of chronic disease. Most importantly, ABHD12 gene therapy reduces macrophage activation and preserves motor function in the sarmopathy model. Thus, we reveal PS as a central component of the SARM1 non-cell autonomous degenerative pathway and develop a novel therapeutic immunomodulator relevant to diseases of PS dysregulation such as PHARC and the many neurodegenerative conditions that feature chronic SARM1 activation.

NMNAT2 is produced in the soma and undergoes anterograde transport into the axon where it functions as an NAD+ biosynthetic enzyme^10^. NMNAT2 is highly labile, and blockade of axon transport via direct injury such as axotomy leads to rapid loss of NMNAT2 from the distal axon^11^. Turnover of NMNAT2 induces a rise in the NMN/NAD+ ratio triggering SARM1 activation and axon degeneration. We first developed an in vitro model of pure SARM1 activation, termed sarmopathy, to investigate the subcellular changes that occur in axons upon genetic versus traumatic loss of NMNAT2. We confirmed that the absence of NMNAT2 was sufficient to trigger full activation of SARM1 and decrease axonal NAD+ but were surprised that this did not also induce axon fragmentation. Rather, sarmopathic axons display blebs and externalized phosphatidylserine while maintaining overall axon integrity. This finding contradicts prior work reporting that acute *Nmnat2* knockdown triggers spontaneous axon degeneration^11^. Thus, the persistence of sarmopathic dKO axons despite ongoing full SARM1 activation suggests the presence of a previously unappreciated axon survival mechanism.. First, dKO axons may develop a compensatory strategy for maintaining axon integrity despite low NAD+ conditions that enhances their resilience to SARM1-mediated metabolic crisis. Indeed, in support of this hypothesis, we and others have reported that mice expressing hypomorphic *Nmnat2* alleles develop and survive in the presence of sub-threshold levels of NMNAT2 that would be expected to trigger spontaneous axon degeneration^8,45,66–68^. Moreover, RNA interference-mediated acute genetic loss of *Nmnat2* may drive spontaneous degeneration due to the inherent additional cellular stress of this technique (e.g. viral infection) that synergizes with SARM1 activation. Loss of a second axon maintenance pathway and/or compensatory mechanism or activation of an additional pro-degenerative pathway may underly our striking finding that axotomy triggers degeneration of sarmopathic dKO axons that would otherwise persist if left unperturbed. Thus, undiscovered factors must be required to complete axon dismantling when SARM1 is activated in the context of developmental *Nmnat2* deletion, and whose discovery could lead to the development of novel axon resiliency strategies.

The effect of ABHD12 upon sarmopathy suggests that dysregulation of PS metabolism is critical to the transcellular axon injury response. ABHD12 regulates lysoPS and oxidized PS (oxPS) species^31^. Altered cellular bioenergetics can lead to excessive production of reactive oxygen species (ROS) and this surplus ROS reacts with lipid membranes generating oxPS^52^ which is readily externalized to the outer plasma membrane leaflet where it acts as an immunomodulatory signal^29,30^. ABHD12 hydrolyzes flipped oxPS via the enzyme’s exofacially-oriented active site^69–71^, reducing excess oxPS. Though we were unable to measure oxPS levels in this study, we observed robust total PS exposure on sarmopathic axons using Annexin V, which indiscriminately binds PS species^28^. Further studies will be required to determine the relative contribution of oxPS to sarmopathic activation of nerve macrophages.

Our prior work demonstrated that macrophages are involved in SARM1-dependent axon loss both at pre-symptomatic and symptomatic disease stages. When and to what degree PS exposure drives pathology throughout disease remains to be elucidated. Likewise, phagocytosis is classically thought of as an immunologically silent program for cell clearance and blocking this process at critical disease stages may lead to nerve inflammation as a result of secondary necrosis or accumulating axonal debris. Therefore, future studies will be needed to dissect the potential pitfalls of chronic oxPS dampening in progressive neurological disease and to understand at which stages of disease pathologic PS exposure drives non-cell-autonomous axon loss amenable to therapy.

Our understanding of PS functions in the nervous system remains incomplete, but several important observations have been made in the context of specific neurodegenerative conditions. AD patients have altered plasma membrane asymmetry^72^ and living neurons with tau filaments aberrantly expose phosphatidylserine and are phagocytosed by microglia^73–75^. Furthermore, phagocytic blockade preserves tau inclusion-bearing neurons, suggesting that phagocytic clearance drives loss of viable neurons. Importantly, the results presented here support our previous findings that SARM1-mediated axon degeneration is not exclusively axon-autonomous and that macrophages respond to early signals from stressed-but-viable axons. This suggests the potential therapeutic value of targeting both PS and SARM1 to treat diseases including SARM1-dependent neuropathies.

## Materials and Methods

### Mouse dorsal root ganglia cultures

Embryos were dissected from embryonic day 13.5 (E13.5) CD-1 (Charles River), E13.5 SARM1 KO, or E13.5 NMNAT2; SARM1 dKO mouse embryos in Dulbecco’s modified Eagle’s medium. DRGs were dissociated in 0.05% trypsin-EDTA (Gibco) at 37°C for 15 minutes, then resuspended in complete medium [Neurobasal E (Gibco) containing 2% B27 (Invitrogen), 100ng/mL nerve growth factor (2.5S Harlan Laboratories), 1mM 5-fluoro-2’-deoxyuridine (Sigma), 1mM uridine (Sigma), and penicillin/streptomycin]. Cell suspensions (1ml/96-well, 10ml/24-well) were placed in the center of wells from 96-or 24-well plates coated with 0.1mg/mL poly-D-lysine (Sigma) and 3mg/mL laminin (Invitrogen) as previously described. After cell attachment, complete medium (100ml/96-well and 500ml/24-well) was added. On DIV5, lentiviruses were added, and axon degeneration assays or metabolite extraction began on DIV6-9.

### Lentivirus transduction

Lentiviruses were produced in HEK293T cells as previously described (Osterloh et al. 2012). Cells were seeded in 6-well plates the day before transfection. FCIV lentivirus vectors harboring cDNAs including SARM1, NMNAT2, and SARM1_E642A_ were co-transfected with pspAx2 (1.2mg) and VSV-G (400ng). Virus-containing culture media were isolated 3 days after transfection and centrifuged for 1 minute at 500 x *g*. Viral supernatant was collected and stored at -80°C. The infection efficiency of all viruses in DRG neurons was assessed by monitoring Venus expression.

### Axon degeneration quantification

Axon degeneration was quantified using ImageJ (NIH) software with a Degeneration Index (DI) algorithm which computes an unbiased degeneration score ranging from 0.0-1.0, with 1.0 representing total fragmentation^76^. Within each well, 6 to 9 distal fields were imaged. Each technical replicate included 3-4 wells per condition and experiments were repeated for at least three independent biological replicates.

### Annexin quantification

AnnexinV fluorescent intensity was quantified using ImageJ. Axonal area was quantified using composite images of bright field and fluorescence from 9 random fields of each well of a 24 well plate. Area corresponding to axon were determined by the bright field image as described previously^76^ and the fluorescent intensity from the axon were quantified. Fluorescence intensity from >40-pixel Annexin positive structures was divided by total axon area. Measurements were performed in triplicate and averaged over three independent experiments.

### Vacor and DO264 treatment

Vacor and DO264 are resolved in DMSO at concentrations of 50mM and 10mM respectively and diluted with the culture medium as indicated. DRG neurons cultured in 24 well were incubated with Vacor or DO264 at the concentration and time as indicated and metabolites were extracted.

### Metabolite collection and measurement

At DIV8, 24-well DRG drop culture medium was replaced with ice-cold 0.9% NaCl solution with or without axotomy, and metabolites were extracted using ice-cold 50% MeOH/water on ice. For axonal metabolite collection, DRG cell bodies were removed immediately after axotomy prior to metabolite collection. Soluble metabolites were further extracted with chloroform and the aqueous phase was lyophilized, then stored at -20°C until LC-MS analysis.

For LC-MS/MS, the metabolite samples were reconstituted with 5mM ammonium formate, centrifuged 12,000 x *g* for 10 minutes, and the cleared supernatant was applied to the LC-MS/MS for metabolite identification and quantification. Liquid chromatography was performed by HPLC system (1290; Agilent) with Synergi Fusion-RP (4.6 3 150mm, 4 mm; Phenomenex) column. Samples (10 ml) were injected at a flow rate of 0.55 ml/min with 5 mM ammonium formate for mobile phase A and 100% methanol for mobile phase B and metabolites were eluted with gradients of 0–7 min, 0%–70% B; 7-8 min, 70% B; 9-12 min, 0% B. The metabolites were detected with Triple Quad mass spectrometer (6460 MassHunter; Agilent) under positive ESI multiple reaction monitoring (MRM). Metabolites were quantified by MassHunter quantitative analysis tool (Agilent) with standard curves. Standard curves for each compound were generated by analyzing NAD+, cADPR, and ATP reconstituted in 5mM ammonium formate.

For NAD+ consumption and synthesis measurements, DRG neurons were incubated with d4-Nam (300uM: 2,3,4,5 deuterium Nam) for 0 or 4 hours and axonal metabolites are collected as described above. Heavy labelled or non-labeled (light) NAD+ were quantified by LC-MS/MS and the rate of consumption was calculated from the % increase or decrease per hour after d4-Nam application.

### LysoPS measurements

LysoPS extraction from cultured DRGs was performed as previously described^77^. Briefly, medium was removed from DRGs cultured in 24 well plates and replaced with 150μl acidic methanol then manually scraped off the plate by pipet. Cell suspensions were transferred to a 1.5ml tube. The obtained mixtures were homogenized by sonication. After initial centrifugation at 1,000 g for 10 min at 4°C, the supernatant was transferred to new tubes and centrifuged at 21,500 g for 10 min at 4°C. 100μl of cleared supernatant was transferred to new tubes, and 10μl of the supernatant was subjected to LC-MS/MS.

### Live DRG imaging

DRG neurons were transduced with lentivirus on DIV 5. Annexin V (#A23204, Thermo Fisher Scientific) was applied according to product instructions.

### AAV constructs and virus injections

AAV constructs were created as previously described^78^. Briefly, AAV8-hSYN-ABHD12-EGFP was generated by the viral vector core of the Hope Center for Neurological Disorders at Washington University in St. Louis. Under light anesthesia with Avertin, 1.3 × 10^13^ viral genomes were injected intrathecally at L6/S1. Viral expression was confirmed by detecting EGFP expression via immunohistochemical analysis of spinal cords.

### Nerve immunohistochemistry

Six μm–thick sections of sciatic nerves were prepared on a cryostat (Leica CM1860), mounted onto slides, and processed as previously described^35^ using the following antibodies: (a) for *activated macrophages*, primary Ab CD68 (1:100, Bio-Rad, MCA1957GA) and secondary Ab anti-rat Cy3 (1:500, Jackson Immunoresearch, 112-165-143); (b) for *total macrophages*, primary Ab Iba-1 (1:500, Wako Chemicals, 019-19741) and secondary Ab anti-rabbit Alexa Fluor 488 (1:500, Invitrogen, A11034); Sciatic nerves were imaged with the Leica DMI 4000B confocal microscope at ×20 magnification.

### Nerve structural analysis

Femoral nerves were processed as previously described^35,79^. Nerves were embedded so that the most distal portion was sectioned. For the light microscope analysis of 400–600nm semithin sections were cut using Leica EM UC7 Ultramicrotome and placed onto microscopy slides. Toluidine blue staining and quantification were performed as previously described^79^. All quantifications were performed by an individual blinded to genotype.

### Inverted screen test

The inverted screen assay was performed as previously described with minor modifications^80^. Mice were placed on a wire mesh screen. Latency to fall for each mouse was recorded, and each mouse underwent 3 trials with 5-minute rest periods. If a mouse did not fall off the mesh screen within 120 seconds, then that time was recorded, and the mouse was taken off the screen.

### Statistics

Unless otherwise stated, data are reported as mean ± SEM. Between-group comparisons were made with 1-way and 2-way ANOVA with post hoc Holm-Sidák multiple comparison test or paired and unpaired 2-tailed *t* tests, as appropriate. 2-sided significance tests were used throughout and *P* < 0.05 was considered statistically significant. All statistics were calculated with the aid of GraphPad Prism 9 (https://www.graphpad.com/scientific-software/prism/) software.

### Study approval

Mice were housed and used following institutional animal study guidelines at Washington University in St. Louis. These protocols received approval from the Washington University IACUC.

## Author Contributions

CBD, YS, AD, and JM conceived the overall study. All authors contributed to the study design. CBD and YS performed the experiments and analyzed the data. DWS generated the *Nmnat2/Sarm1* dKO mouse and provided intellectual input. AS assisted with animal husbandry and histological analyses. CBD wrote the manuscript and prepared all figures. AJB, YS, JM, and AD oversaw the analyses and revised the manuscript. All authors gave final approval of the manuscript.

## Acknowledgments

We thank Milbrandt and DiAntonio labs members for technical support and critical feedback on this manuscript.

## Funding

This work was supported by National Institutes of Health grants (R01NS087632 and R01NS133348 to JM and AD, R01NS119812 to JM, AD, and AJB, and R01AG013730 to JM).

## Declaration of interests

AD and JM are cofounders, scientific advisory board members, and shareholders of Disarm Therapeutics, a wholly owned subsidiary of Eli Lilly.

